# Clinical Significance of the Stromatic Component in Ovarian Cancer: Quantity Over Quality in Outcome Prediction

**DOI:** 10.1101/2023.06.27.546712

**Authors:** Emil Lou, Valentino Clemente, Marcel Grube, Axel Svedbom, Andrew Nelson, Freya Blome, Annette Staebler, Stefan Kommoss, Martina Bazzaro

## Abstract

**Background:** The tumor stroma is composed of a complex network of non-cancerous cells and extracellular matrix elements that collectively are crucial for cancer progression and treatment response. Within the realm of ovarian cancer, the expression of the stromal gene cluster has been linked to poorer progression-free and overall survival rates. However, in the age of precision medicine and genome sequencing, the notion that the simple measurement of tumor-stroma proportion alone can serve as a biomarker for clinical outcome is a topic that continues to generate controversy and provoke discussion. Our current study reveals that it is the quantity of stroma, rather than its quality, that serves as a clinically significant indicator of patient outcome in ovarian cancer.

**Methods:** This study leveraged the High-Grade-Serous-Carcinoma (HGSC) cohort of the publicly accessible Cancer Genome Atlas Program (TCGA) along with an independent cohort comprising HGSC clinical specimens in diagnostic and Tissue Microarray formats. Our objective was to investigate the correlation between the Tumor-Stroma-Proportion (TSP) and progression-free survival (PFS), overall survival (OS), and response to chemotherapy. We assessed these associations using H&E-stained slides and tissue microarrays. Our analysis employed semi-parametric models that accounted for age, metastases, and residual disease as controlling factors.

**Results:** We found that high TSP (>50% stroma) was associated with significantly shorter progression-free survival (PFS) (p=0.016) and overall survival (OS) (p=0.006). Tumors from patients with chemoresistant tumors were twice as likely to have high TSP as compared to tumors from chemosensitive patients (p=0.012). In tissue microarrays, high TSP was again associated with significantly shorter PFS (p=0.044) and OS (p=0.0001), further confirming our findings. The Area Under the ROC curve for the model predicting platinum was estimated at 0.7644.

**Conclusions:** In HGSC, TSP was a consistent and reproducible marker of clinical outcome measures, including PFS, OS, and platinum chemoresistance. Assessment of TSP as a predictive biomarker that can be easily implemented and integrated into prospective clinical trial design and adapted to identify, at time of initial diagnosis, patients who are least likely to benefit long-term from conventional platinum-based cytotoxic chemotherapy treatment.

## Introduction

The role of the tumor microenvironment (TME) in predicting invasive behavior, metastatic potential, and chemoresistance of tumors is an emerging field at the intersection of tumor biology and clinical care^1–5^. Tumor-stroma proportion (TSP) is an assessment of the extent of involvement of tumor with stromatous TME components and has been associated with poor prognosis in a broad range of cancers^6–8^. Ovarian cancer is a heterogeneous and potentially invasive form of carcinomas. Nearly 1 out of every 3 patients develops resistance to platinum-based chemotherapy, which remains the backbone of standard-of-care treatment for advanced-stage forms of this disease^9, 10^.

New therapeutic targets for the treatment of recurrent ovarian cancer are urgently needed, especially for these patients with disease that is refractory to standard-of-care platinum chemotherapy. Even in the era of molecular oncology and advances in our understanding of the genomic drivers and passengers implicated in the carcinogenesis and metastatic potential of ovarian carcinomas, there is limited efficacy of molecular therapeutics targeting these mutations. This state of the field provides an opportunity to examine the biology of the ovarian TME as potential alternative targets for treatment. In 2019, we published results from a prospective study of women with newly diagnosed ovarian tumors that showed that tumors harboring a higher tumor-stroma proportion (TSP) predicted eventual emergence of platinum drug resistance^11^. This study was the first study to evaluate TSP from assessment at time of ovarian cancer diagnosis and provide correlation to eventual emergence of drug resistance over time. The patients were monitored over time for response of their tumors to standard-of-care platinum-based chemotherapy, and their cases were designated as platinum-sensitive or platinum-resistant based on standard definitions in the field (recurrence after six months following first-line treatment, vs. recurrence prior to six months following first-line treatment, respectively). TSP assessed from initial biopsy or surgical specimen was determined using slides used for establishing histopathologic diagnosis. The discovery that high TSP was associated with platinum-resistance offered TSP as a potentially effective and low-cost predictive biomarker of drug resistance that, if confirmed and validated, could advance the field of ovarian cancer treatment by helping to tailor treatment for patients in whom platinum therapy was destined to be less effective than in others.

Thus, we next sought to confirm these results in a larger patient cohort. To do this, we accessed the Cancer Genome Atlas Program (TCGA) High-Grade-Serous-Carcinoma (HGSC) cohort and created a larger collaborative network with access to nearly 200 specimens of ovarian carcinoma, linked to clinical outcomes data that included information on progression-free survival, overall survival, and platinum chemotherapy resistance. We excluded intrinsic variability due to different biology of multiple histologic subtypes of ovarian cancer by choosing to use high grade serous carcinomas (HGSC) only. We performed univariate and multivariate analysis of chemoresistance and survival in this cohort and smaller cohorts in order to exclude the effect of confounders and identify specific subpopulations that may benefit the most from the use of TSP as a predictive biomarker. Here, we provide confirmation and validation of our previous finding associating high TSP with chemoresistance using this larger dataset that high TSP is not only predictive of chemotherapy resistance, but also indicative of worse PFS and OS, and thus worsened prognosis, in patients with high grade serous ovarian carcinomas. While our results align with previous studies suggesting that the stromal gene cluster is associated with unfavorable progression-free and overall survival outcomes in ovarian cancer ^12, 13^ our findings further indicate that this relationship is primarily related to the quantity or extent of stromal involvement.

## Materials and Methods

### The Cancer Genome Atlas (TCGA) Cohort

We interrogated the TCGA dataset for cases of ovarian cancer which had both annotated clinical data and hematoxylin and eosin (H&E)-stained that had been used for diagnosis and then scanned into the TCGA archives. This search was performed using the following filters: primary site “ovary” → data type “slide image” → experimental strategy “Diagnostic slide” → disease type “cystic, mucinous and serous neoplasms”. This algorithm retrieved a total of 106 cases. Three patients were excluded due to the absence of clinical data (UUID 01abf2cf-541c-4610-8ce8-318016db6e59) or poor quality of the specimen (UUIDs 700e91bb-d675-41b2-bbbd-935767c7b447 and 8628f1b2-3763-4ba9-a375-083874bb18f2). The final cohort consisted of 103 patients whose cases were diagnosed and treated at multiple institutions between 1993 and 2013. ICD-O-3 codes for topographic origin were C56.9 (malignant neoplasm of the ovary) and C48.0 (malignant neoplasm of retroperitoneum). ICD-O-3 codes for pathological diagnosis were: 8441/3 (Serous cystadenocarcinoma, including serous adenocarcinoma and serous carcinoma), 8440/3 (cystadenocarcinoma) and 8460/3 (serous cystadenocarcinoma). All patients had been treated with surgery followed by standard-of-care adjuvant chemotherapy. No history of neoadjuvant chemotherapy was reported. Demographic information and patient characteristics are reported in Table 1. Representative microphotographs of TSP in the TCGA cohort are given in Figure 1A.

**Figure 1.**
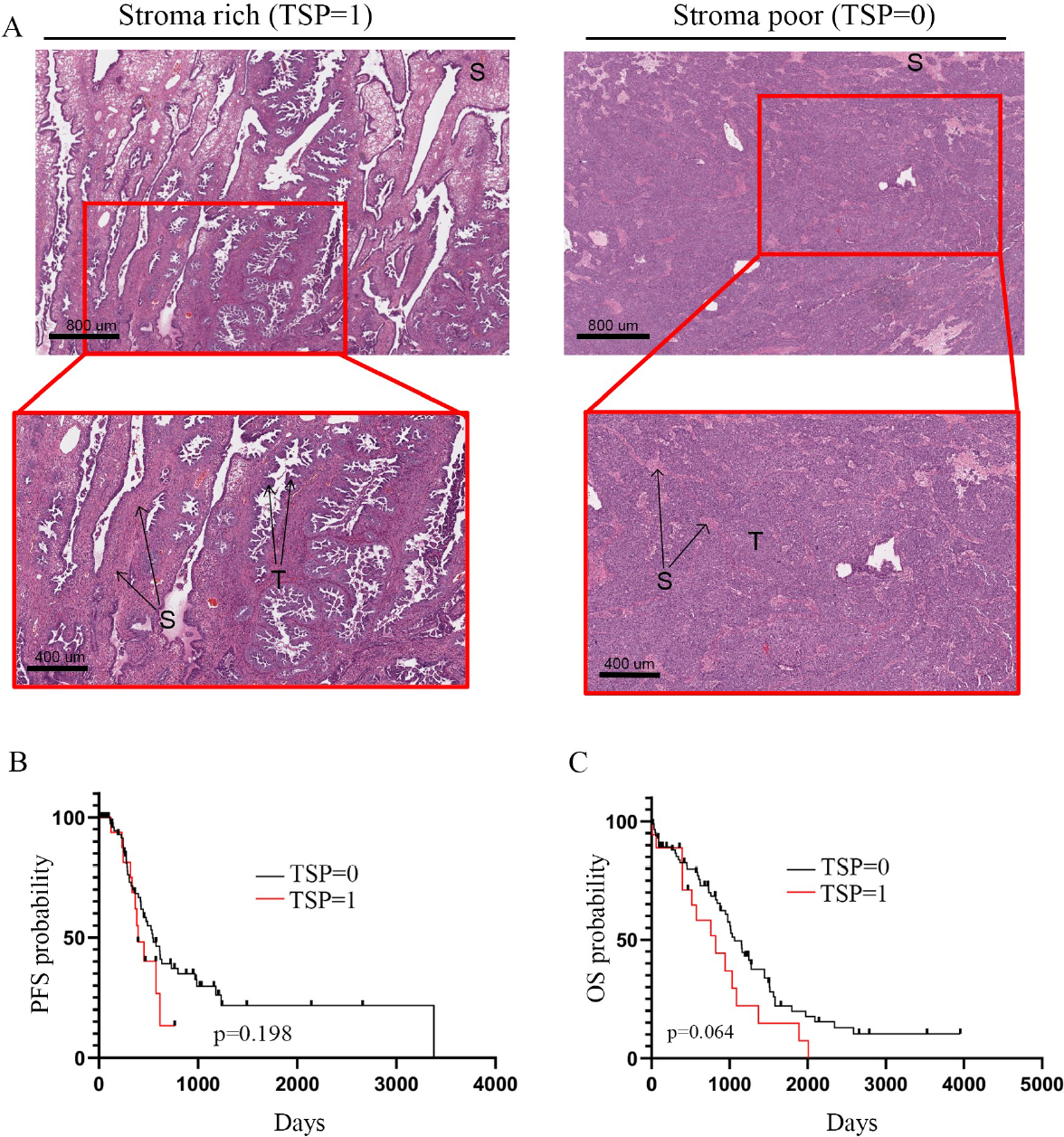
Association and between TSP and clinical outcome in clinical specimens ovarian cancer in the TCGA cohort. **A**. *Left*, representative images of stroma rich (TSP=1) specimens in the TCGA cohort. This corresponds to the slide ID TCGA-23-1024-01Z-00-DX1.B9194D3F-C6F4-4FC8-B0CA-6E347FF4F885 and to the case UUID: 8ac6ce06-ab50-4ab4-a469-9f8bf01be963. *Right,* representative images of stroma poor (TSP=0) clinical specimens. This corresponds to the slide ID: TCGA-23-2079-01Z-00-DX1.01383205-B7A7-4529-9207-880687650A50 and UUID: e3ea5137-76b8-4afc-9b80-ed74714e7172. S=stromal component within the tumor, T=cancerous cells within the specimen. **B**. Kaplan-Meier curve for progression-free survival (PFS) in patients with low TSP (TSP=0) *vs.* high TSP (TSP=1) (p=0.198). **C**. Kaplan-Meier curve for overall survival (OS) in patients with low TSP (TSP=0) *vs.* high TSP (TSP=1) (p=0.064).

**Table 1.** Patient Demographics, Clinical Characteristics, and Tumor Stroma Proportion (TSP) of the 103 Women Diagnosed with Ovarian Carcinoma with data from TCGA. Cases with low TSP (<50%) are labeled as TSP=0, and cases with high TSP (>50%) are labeled as TSP=1.

| Characteristic | Level | All | TSP=0 | TSP=1 | p-value |
| --- | --- | --- | --- | --- | --- |
| **Number of patients** |  | 103 | 85 | 18 |  |
| **Age at diagnosis, mean (SD)** |  | 61.6 (11.1) | 61.4 (11.2) | 62.7 (11.1) | 0.66 |
| **FIGO stage** | Stage IC | 2 (1.9%) | 2 (2%) | 0 (0%) | 0.91 |
|  | Stage IIA | 1 (1.0%) | 1 (1%) | 0 (0%) |  |
|  | Stage IIB | 1 (1.0%) | 1 (1%) | 0 (0%) |  |
|  | Stage IIC | 2 (1.9%) | 1 (1%) | 1 (6%) |  |
|  | Stage IIIB | 1 (1.0%) | 1 (1%) | 0 (0%) |  |
|  | Stage IIIC | 72 (69.9%) | 59 (69%) | 13 (72%) |  |
|  | Stage IV | 23 (22.3%) | 19 (22%) | 4 (22%) |  |
|  | Unknown | 1 (1.0%) | 1 (1%) | 0 (0%) |  |
| **Residual disease** | No | 21 (20.4%) | 18 (21%) | 3 (17%) | 0.76 |
|  | Yes | 76 (73.8%) | 63 (74%) | 13 (72%) |  |
|  | Unknown | 6 (5.8%) | 4 (5%) | 2 (11%) |  |
| **Chemoresistance** | No | 60 (58.2%) | 49 (57.6%) | 12 (66.7%) | 0.78 |
|  | Yes | 12 (11.7%) | 12 (14.1%) | 2 (11.1%) |  |
|  | Unknown | 31 (30.1%) | 24 (28.2%) | 4 (22.2%) |  |

### Human Subjects Tübingen Cohort

The Tübingen cohort consisted of 192 clinical specimens derived from patients assessed at Tübingen University Hospital in Tübingen, Germany from 2004 to 2014 for suspected or known ovarian carcinomas, and who underwent debulking surgical resection of tumors followed by standard-of-care systemic therapy. Archival tissues were used with approval from the Internal Ethics Review Board (Ethikkommission) of the Medical Faculty of the University of Tübingen (Project-Nr: 645/2012 BO2). Demographic information and patient characteristics are reported in **Supplementary Table 1**.

### TSP scoring

The scoring was performed by one clinician (V.C.) and one pathologist (A.N.) who examined (10X) the hematoxylin/eosin (H&E)-stained slides to identify representative sections of the primary tumor. For scoring of the 192 whole sections: after identifying the most densely populated cellular areas of the slide, the relative amounts of tissue represented by cancer cells vs. stromal tissue were identified and labeled using 50% as a previously validated cutoff^11^. Tumors with a stroma proportion <50% was classified as low (designated as tumor-stroma ratio (TSP)=0 or low TSP) and tumors with a stroma proportion ≥ 50% as high TSP (TSP=1 or high TSP). Stromal tissue that was not in clear continuity with the peritumoral stroma was not included in the evaluation. Necrotic, mucinous, empty, or large inflammatory areas were excluded from the scoring.

For scoring of the Tissue Microarray (TMA): of the 192 clinical specimens in the Tübingen cohort, 185 were available in the TMA format. These TMAs consisted of 6 punches for each of the 185 clinical specimens built using six 0.8mm diameter cores for each tumor. A total of 1110 (185X6) H&E stained were identified and labeled using 50% as the cutoff point. A single TSP category was assigned to each patient based on the mode from the six total TSPs values. Based on this process, of the total 185 patients, 143 were labeled as stroma-poor (low TSP, TSP=0) and 42 were labeled as stroma-rich (high TSP, TSP=1), corresponding to a 9.2% discrepancy between diagnostic slide and TMA-based scoring. Finally, univariate analysis of the outcome was performed using standard Kaplan-Meier survival curves and log-rank tests to compare progression-free and overall survival TSP=1 and TSP=0 patients.

### Covariates

In multivariate analysis we controlled for age at diagnosis, residual disease, primary metastases (yes/no), and lymph node spread (yes). Chemoresistance was defined as progressive disease during chemotherapy or less than 180 days after the first adjuvant regimen. Residual disease was reported as: a) No residual disease, b) Residual disease 0 = <1 cm remaining (following suboptimal debulking), or c) Residual disease 1 = >1 cm (optimal debulking). All covariates were measured at baseline.

### Statistical analysis

We presented categorical variables using frequency and percentage, and continuous variables using mean and standard deviation or median and interquartile range depending on the approximately normality of the underlying distributions. We compared categorical variables using chi-square tests and continuous variables using t-tests by virtue of the central limit theorem. In time-to-event analyses, patients were followed until first of failure or right-censoring, defined as loss to follow-up or date of data extraction. Kaplan-Meier time-to-event functions ^14^ were fit to estimate progression free and overall survival and we fit Cox-Proportional Hazards (PH) models ^15^ to address confounding factors. We tested for differences in time-to-events between groups using the Wald-statistic for the hazard ratio from Cox PH models and validated the proportionality assumption by inspecting the Schoenfeld residuals^16^. The proportionality assumption was valid unless otherwise noted. We fit a logistic regression model to estimate the association between platinum resistance and TSP controlling for the covariates (age at diagnosis, lymph node spread (yes/no), distant metastasis (yes/no), and residual disease (yes/no).

## Results

### Patient demographics and TSP assessment in the TCGA cohort

In 2019, we reported our finding from a small prospective cohort of twenty-four patients that high TSP is associated with platinum chemoresistance in ovarian cancer^11^. To add supporting evidence that high TSP correlates with chemoresistance, and to also investigate potential concurrent association with prognosis, we first used a retrospective dataset available via TCGA consisting of H&E digitized slides used for diagnostic purposes and annotated with clinical outcomes. This TCGA cohort consisted of 103 patients with a histopathologic diagnosis of ovarian carcinomas. Patient demographics and clinical characteristics, including FIGO stage, residual disease following surgery, and chemoresistance, are listed in Table 1. The average age of patients at the time of diagnosis was 61.6 years. Most patients (69.9%) were FIGO stage 3; slightly less than 2/3 had residual disease following surgical debulking. Although only 11.7% of all cases had chemoresistant disease, the 30.1% of cases had unknown chemoresistance status thus limiting ability to provide confirmation in this analysis. We determined TSP using the digitized images using our previously reported cut-off of TSP <50% and >50%^11^ and designated the ratios as Tumor-Stroma Ratio (TSP)=0 (TSP-low) or TSP=1 (TSP-high), respectively. Representative images of H&E-stained slides of TSP=0 and TSP=1 are shown in Figure 1A.

### Characteristics of patients with high vs. low tumor-stroma proportions in the TCGA cohort

We compared patient demographic characteristics, including age at diagnosis, stage, presence of residual disease, and platinum-resistant vs. –sensitive form of the disease in relation to TSP (Table 1). Of the total 103 patients in the TCGA dataset, 18 patients (17.5%) had TSP=1 (high TSP), and the remaining 85 patients had tumors of TSP=0 (low TSP). No significant differences were found between TSP=0 and 1 tumors when comparing clinical and demographic characteristics.

### High TSP trends towards lower progression-free and overall survival in the TCGA cohort

We conducted a correlation analysis between TSP and outcome measures including progression-free survival and overall survival in the TCGA cohort. Our analysis revealed that, while not statistically significant, a TSP=1 status appeared to be associated with worse progression-free survival several years following initial diagnosis and treatment (p=0.198) (Figure 1B). Additionally, the association between TSP=1 and overall survival trended toward but did not reach statistical significance (p=0.064), again following a similar pattern that differentiated the two groups after the first several years from diagnosis. (Figure 1C). We observed no significant correlation between TSP levels and chemoresistance in the TCGA cohort (Table 1); as noted above, chemoresistance status was unknown or otherwise not listed in approximately 30% of all cases in this dataset. Thus, overall, this initial analysis provided a potential signal that high TSP could be associated with worse PFS and OS, but further investigation would be needed using a larger and more complete dataset.

### Patient demographics and TSP assessment in the Tübingen cohort

Having previously demonstrated a significant association of TSP with ovarian cancer chemoresistance in a small prospective cohort, but no statistically significant association using the larger but limited TCGA dataset, we next sought confirmation using an even larger and more comprehensively annotated cohort available at the University of Tübingen in Germany. We performed retrospective analysis of this 192-patient cohort of patients diagnosed with high-grade serous carcinoma (HGSC) of the ovaries; hereafter we will refer to this dataset as the ‘Tübingen cohort’. Patient demographics and clinical characteristics, including FIGO stage and presence of residual disease following surgery, are listed in **Supplementary Table 1**. The average of patients at time of diagnosis was 63.7 years. The majority of patients (69.8%) were FIGO stage 3, with half of patients having node-positive disease. More than 2/3 of patients had no evidence of distant metastasis. Over half of patients (56.3%) had residual disease following surgical debulking. Similar to what we did for the TCGA cohort and in our previously published work^11^ here too we assessed TSP using standard H&E-stained histologic specimens used for confirmation of histologic diagnosis and again designated the ratios as TSP=0 (TSP-low) or TSP =1 (TSP-high), respectively. Representative images of H&E-stained slides from the Tübingen cohort of TSP-high and TSP-low cases are provided in Figure 2A.

**Figure 2.**
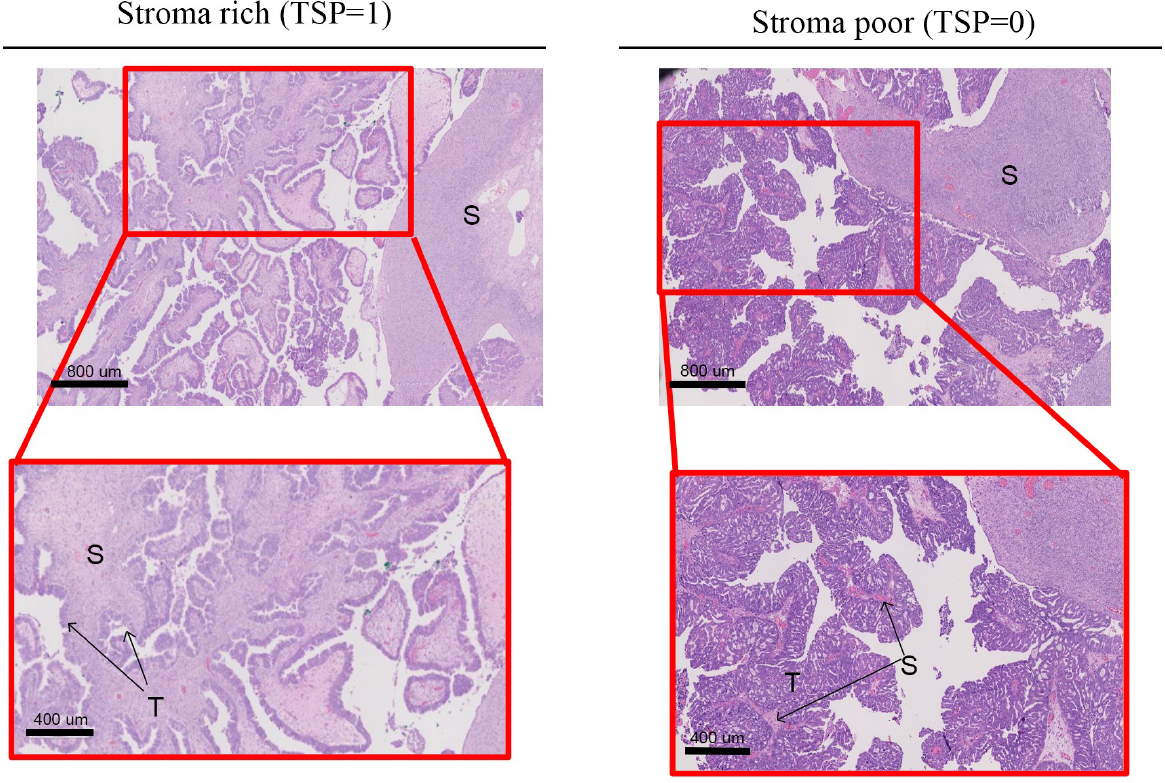
A. Stroma rich and poor clinical specimens of High-Grade-Serous-Carcinoma (HGSC) of the ovaries in the Tübingen cohort and PFS and OS. **A.** *Left,* representative images of stroma rich (TSP=1) specimens in the Tübingen cohort. *Right*, representative images of stroma poor (TSP=0) specimens in the Tübingen cohort. S=stromal component within the tumor, T=cancerous cells within the specimen. **B**. Kaplan-Meier curve for progression-free survival (PFS) in patients with low TSP (TSP=0) *vs.* high TSP (TSP=1) (p=0.016). **C**. Kaplan-Meier curve for overall survival (OS) in patients with low TSP (TSP=0) *vs.* high TSP (TSP=1) (p=0.006).

### Characteristics of patients with high vs. low tumor-stroma proportions in the Tübingen cohort

We compared patient demographic characteristics including age at diagnosis, stage, presence of residual disease, and platinum-resistant vs. –sensitive form of disease in relation to TSP (Table 2). Of the total 192 patients in the dataset, 41 patients (21.4%) had TSP=1 (high TSP), and the remaining 151 patients had tumors of TSP=0 (low TSP). Patients with high TSP had a higher rate of FIGO stage IV cases at time of initial diagnosis (p=0.037). As the most frequent T and N stages in both groups were T3c and N0, respectively, this is likely explained by the higher rate of distant metastases in TSP=1 patients (p=0.012). There were no other significant differences in age, tumor size, number and extent of affected lymph nodes between patients with TSP 0 *vs*. 1.

**Table 2.**
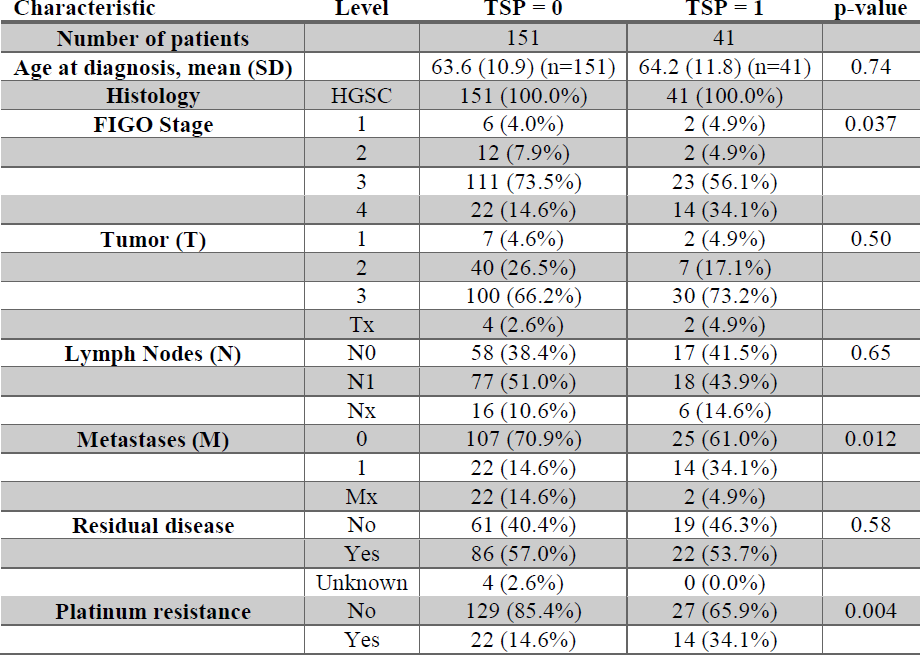
Patient Demographic and Clinical Characteristics of patients divided by Tumor Stroma Proportion (TSP)

| Characteristic | Level | TSP = 0 | TSP = 1 | p-value |
| --- | --- | --- | --- | --- |
| **Number of patients** |  | 151 | 41 |  |
| **Age at diagnosis, mean (SD)** |  | 63.6 (10.9) (n=151) | 64.2 (11.8) (n=41) | 0.74 |
| **Histology** | HGSC | 151 (100.0%) | 41 (100.0%) |  |
| **FIGO Stage** | 1 | 6 (4.0%) | 2 (4.9%) | 0.037 |
|  | 2 | 12 (7.9%) | 2 (4.9%) |  |
|  | 3 | 111 (73.5%) | 23 (56.1%) |  |
|  | 4 | 22 (14.6%) | 14 (34.1%) |  |
| **Tumor (T)** | 1 | 7 (4.6%) | 2 (4.9%) | 0.50 |
|  | 2 | 40 (26.5%) | 7 (17.1%) |  |
|  | 3 | 100 (66.2%) | 30 (73.2%) |  |
|  | Tx | 4 (2.6%) | 2 (4.9%) |  |
| **Lymph Nodes (N)** | N0 | 58 (38.4%) | 17 (41.5%) | 0.65 |
|  | N1 | 77 (51.0%) | 18 (43.9%) |  |
|  | Nx | 16 (10.6%) | 6 (14.6%) |  |
| **Metastases (M)** | 0 | 107 (70.9%) | 25 (61.0%) | 0.012 |
|  | 1 | 22 (14.6%) | 14 (34.1%) |  |
|  | Mx | 22 (14.6%) | 2 (4.9%) |  |
| **Residual disease** | No | 61 (40.4%) | 19 (46.3%) | 0.58 |
|  | Yes | 86 (57.0%) | 22 (53.7%) |  |
|  | Unknown | 4 (2.6%) | 0 (0.0%) |  |
| **Platinum resistance** | No | 129 (85.4%) | 27 (65.9%) | 0.004 |
|  | Yes | 22 (14.6%) | 14 (34.1%) |  |

Approximately 25-30% of patients with HGSOC develop platinum-resistant tumors. We have previously reported that high TSP is predictive of platinum resistance, thus in this cohort we compared TSP=0 *vs.* TSP=1 in relation to response to platinum-based chemotherapy. There was a significantly higher proportion of TSP=1 cases that were platinum-resistant (34.1%) as compared to TSP=0 cases (14.6%) (P=0.004) (Table 2). In contrast, more TSP=0 cases were platinum-sensitive (85.4%) than was seen in TSP=1 cases (65.9%).

### High TSP is associated with lower progression free and overall survival in all patients with HGSOC in the Tübingen cohort

Among all patients with HGSOC, patients with TSP=0 had a higher probability of progression-free-survival (PFS) than patients with high TSP=1 at all timepoints during follow-up (p=0.016) (Figure 2B). This was also true when we adjusted for age at diagnosis, primary metastasis, and residual disease, at which time TSP=1 was significantly (p=0.015) associated with lower PFS (HR: 1.587; 95% CI: 1.093-2.302) (**Supplementary Table 2)**. The presence of residual disease after surgery – an established prognostic factor for ovarian cancer -- was also associated with worse outcome. However, the presence of distant metastases, which was more common in the TSP=1 group, was not associated with PFS (p=0.529).

In terms of Overall Survival (OS) patients with TSP=0 also had a higher probability of OS at all time points except for the initial months (p=0.006) (Figure 2C). When we adjusted for age at diagnosis, primary metastasis and residual disease, TSP=1 was significantly (p=0.002) associated with lower OS (HR: 1.586; 95% CI: 1.093-2.302) (Table 3). As seen with PFS, the presence of residual disease was associated with shorter OS, together with metastases and age.

**Table 3.** Multivariate analysis of overall survival.

| Variable | Haz. ratio | P-value | [95% conf. | interval] |
| --- | --- | --- | --- | --- |
| TSP | 2 | 0.002 | 1 | 3 |
| Age | 1 | <0.001 | 1.018 | 1 |
| Metastasis | 2 | 0.032 | 1.04 | 2 |
| Residual Disease | 3 | <0.001 | 2 | 4 |

The above assessments of PFS and OS using the Tübingen cohort were performed using a whole-slide approach of the entirety of tumor specimen on each slide. Many studies retrospectively assessing reactive tumor stroma patterns utilize an approach based on tissue microarrays (TMA) constructed using available resection specimens. As this approach is commonly used, we wanted to confirm whether the TMA approach would yield the same results as whole-slide assessment. Of the 192 clinical specimens in the Tübingen cohort, 185 were available in the TMA format for this next analysis. As described in detail in Materials and Methods, these TMAs consisted of 6 punches for each of the 185 clinical specimens built using six 0.8mm diameter cores for each tumor; thus this difference in tumor preparation could account for potential differences in outcomes. Representative images of the TMA sections are shown in Figure 3A. Of these 185 TMA-constructs, we designated 143 were labeled as low TSP, and 42 were labeled as high TSP. These numbers presented a 9.2% discrepancy between the whole-slide H&E-stained diagnostic slides presented above and the TMA-based coring. Nonetheless, univariate analysis of these TMA-constructs again confirmed a statistically significant association of high TSP with lower PFS and OS in the Tübingen cohort (Figure 3 B-C).

**Figure 3.**
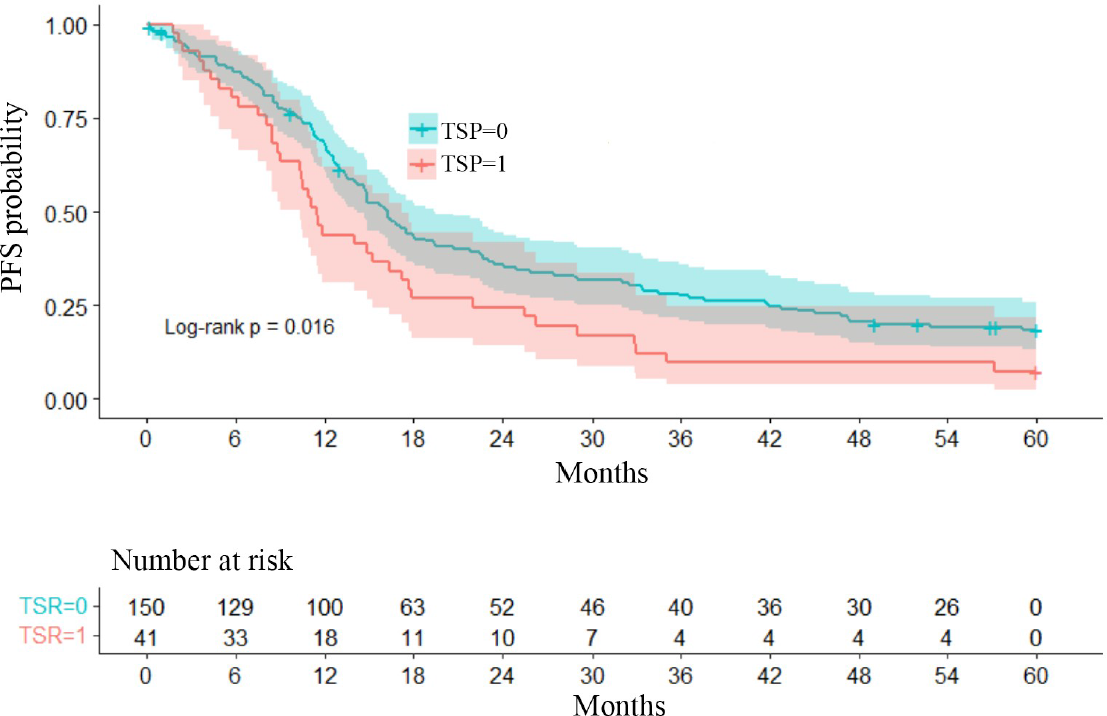
Validation of the use of TMAs to predict overall and progression-free survival based on TSP in the Tübingen cohort. **A.** *Top,* Representative images of stroma rich. **B.** Kaplan-Meier curve for progression-free survival (PFS) in patients with low TSP (TSP=0) vs. high TSP (TSP=1) (p=0.0449; HR 1.675 95% CI 1.012-2.772) as defined by scoring TMAs. **B.** Kaplan-Meier curve for overall survival (OS) in patients with low TSP (TSP=0) vs. high TSP (TSP=1) as defined by scoring TMAs (p=0.0001; HR 2.491 95% CI 1.585-3.912).

### Confirmation of the Association of TSP with platinum chemoresistance in patients with HGSC in the Tübingen cohort

Finally, we used the Tübingen cohort to validate our previously reported findings from 2019 that high TSP is indeed associated with emerging of platinum chemotherapy resistance in women with ovarian cancer. We performed multivariate analysis of cases based on chemoresistance and confirmed a statistically significant difference in which TSP-high cases correlated with platinum-chemoresistance (p=0.012; Odds Ratio 2.861) after adjusting for age, primary metastasis and residual disease (Table 4). Furthermore, the Area Under the ROC curve for the model predicting platinum resistance based on age at diagnosis, metastasis, residual disease, lymph node spread, and TSP was estimated at 0.7644 (Figure 4). Taken together, these results not only suggest that chemoresistance in patients with TSP=1 may be responsible for the poorer outcome also observed in these patients, but also suggest that TSP may have specificity and sensitivity high enough to be employed in the clinical setting to predict which patients are going to respond to treatments or benefit from newer therapeutic options.

**Figure 4.**
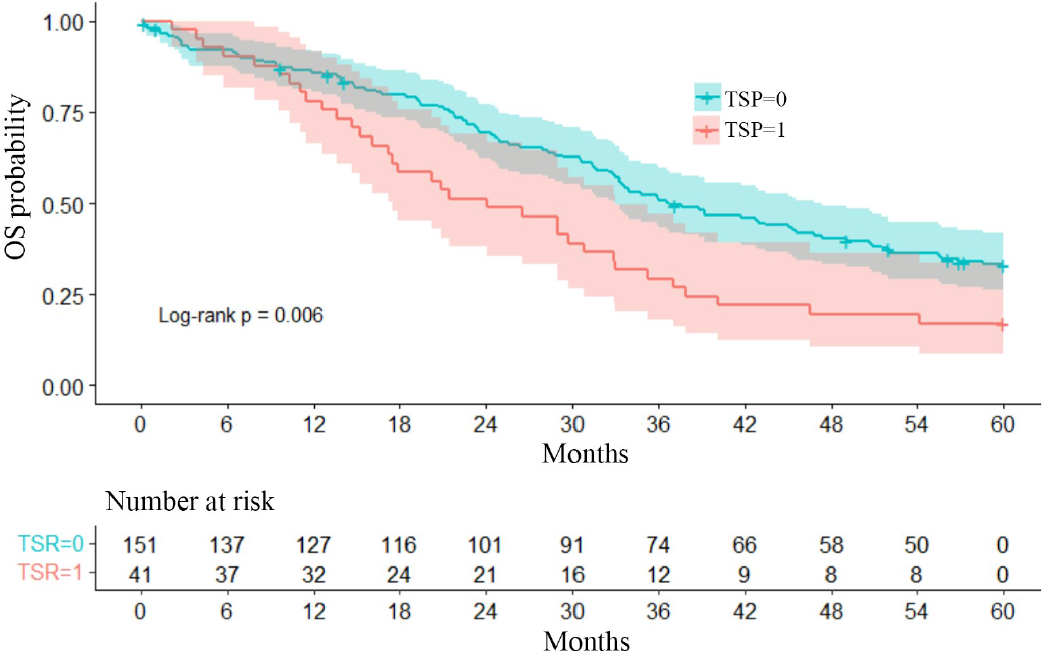
Stroma rich tumors as predictors of platinum resistance in the Tübingen cohort. Receiver operating characteristic (ROC) curves of stroma rich tumors (TSP=1) and risk factors (metastasis, residual disease, lymph node spread) indicating the ability of TSP=1 to differentiate chemoresistant and chemosensitive patients. The diagonal line indicates no predictive value. Area under the ROC curve is 0.7644.

**Table 4.** Multivariate analysis chemoresistance.

| **Variable** | **Odds ratio** | **[95%**<br>**conf.** | **interval]** | **P>t** |
| --- | --- | --- | --- | --- |
| **TSP** | 2.861 | 1.256 | 6,515 | 0.012 |
| **Age at**<br>**diagnosis** | 0.991 | 0.958 | 1.025 | 0.600 |
| **Metastases** | 2,059 | 0.877 | 4.831 | 0.097 |
| **Residual**<br>**disease** | 1.961 | 0.862 | 4.462 | 0.108 |

## Discussion

TSP is a readily assessable property of the tumor microenvironment that holds strong promise not just as an indicator of prognosis, but perhaps more importantly as a predictive biomarker of likely emergence of resistance to standard-of-care chemotherapy over time. TSP has been published widely and defined in previous studies in binary fashion, in other words as tumors with stromatous portion comprising less than or more than 50% stroma^7, 8^. In 2019, we reported our finding from a small prospective cohort of patients that TSP is associated with platinum chemoresistance^11^. Here, we present further supportive evidence that high TSP correlates with platinum chemoresistance and also lower PFS and OS in women with ovarian cancer, using a confirmatory dataset focused on 192 patients with a histopathologic diagnosis of HGSOC.

In the current study, high TSP was the only factor significantly (p=0.012) predictive of emergence of platinum chemotherapy resistance following first-line treatment. The risk was even higher in patients with no metastases at diagnosis. These results not only confirm that higher TSP is associated with development of chemoresistance to current standard of care treatments in a large cohort of HGSC, but also potentially support the notion that this might be the underlying cause of the poorer outcome also observed in these patients. Further studies would be needed to determine factors that drive this role of TSP in chemoresistance at the cellular and molecular level. Our team has investigated molecular and cellular factors that induce and propagate emergence of chemoresistance in patients with ovarian cancer^17–20^, including tumor cell communication networks that bridge malignant cells distributed throughout heterogeneous stroma-rich tumors that present with higher TSP^21^. These communication networks are induced by hypoxia^22^ and other characteristic properties of proliferating tumors^23^ and are just one example of factors that play a role in driving TSP-associated ovarian cancer chemoresistance.

The data presented here provide confirmation of high TSP as a useful companion predictive biomarker for future emergence of platinum resistance of ovarian carcinomas, and also support the notion that it can predict worse prognosis as well. At face value, these results do differ from recently reported results from Micke et al., who examined prognostic impact of tumor stroma fraction in 2,732 cases of resected tumors representing sixteen separate solid tumor types; of this total, there were 197 cases categorized as Ovarian Carcinoma, with an additional 49 cases listed as High Grade Serous Ovarian Cancer (HGSOC)^6^. The investigators utilized TMA-constructs, which the authors acknowledge carry limitations in potential biasing of assessable portions of resected tumors in context of intratumoral heterogeneity. Using deep learning methods, they identified and quantified stroma and analyzed differences using median levels of stroma fraction as the primary cutoff. The median cutoff used for differentiating stroma-rich and stroma-poor fractions was well below 50% for ovarian cancer cases overall (n=150 reported) and HGSOC (n=49)^6^. The authors determined that there was no significant association of TSP with prognosis in many cancer types tested in their study, including both categories of ovarian carcinomas, within the context of reliance on TMAs and the variable cutoff between different tumor types. For our study, as we sought to confirm our prior findings in relation to chemoresistance, and also consistent with other studies using a 50% cutoff in ovarian and other cancers^7, 24–26^, we opted to utilize this 50% value. We also analyzed both TMA and whole-slide sections of surgical specimens to confirm consistency of findings between both preparative techniques, and indeed confirm the findings as described.

We found that TSP was also associated with metastases at diagnosis. There were approximately 65 M1 cases in the dataset, and they included both FIGO stage IVa (ie intraperitoneal mets) and FIGO Stage IVB (extraperitoneal metastases), and were provided together in our clinical dataset unassociated with the site of metastatic disease. It is conceivable that there are differences in platinum resistance between IVA and IVB cases which were not uncovered in this study.

Overall, these data support the notion that higher TSP is at least associated with and may even drive a more aggressive form of HGSC. These results serve as confirmation of our findings in a smaller prospective cohort from which we first reported potential predictive nature of TSP for chemoresistance in ovarian cancer. This suggests not only that high TSP could help identifying patients who may benefit more from alternatives to platinum based regimens, but also that this would be particularly useful for patients with the most common stage at diagnosis in HGSC and, furthermore, who could achieve the best therapeutic results based on chemotherapy.

Limitations of this study include limitations inherent in any retrospective dataset, the relatively inexact nature of assessment of residual disease following attempt at maximal cytoreduction and debulking, and the manual process required to review slides for TSP assessment. In general, some of the subset sample sizes were small when analyses were stratified by metastasis and then further split into exploration and validation datasets. In addition, intratumoral heterogeneity is a known entity that is an inherent factor in any tissue-based evaluation study; whether TSP varies widely enough within each ovarian tumor is a point of speculation and an unknown factor. Furthermore, inter-tumoral heterogeneity of TSP between primary ovarian tumors and metastatic tumors to distant sites, which may be enriched in mesenchymal subpopulations and may have inherently high TSP, remains an additional aspect that can be assessed in future prospective validation studies.

In conclusion, we provide confirmation of high TSP as a predictive biomarker of platinum chemotherapy resistance in ovarian carcinomas. This marker is particularly useful for evaluating prognosis as well as drug resistance in cases of HGSOC that undergo maximal debulking surgery, which is usually associated with best possible prognosis. Here we demonstrate that TSP serves as an additional stratifying factor for determining cases at highest risk of recurrence and death that are likely due in part to lack of efficacy of standard platinum-based chemotherapies. With this confirmation, we propose that TSP should be further standardized and incorporated into prospective clinical trials as a correlative predictive biomarker for drug resistance.

## Acknowledgements

Dr. Emil Lou would like to thank the Minnesota Ovarian Cancer Alliance and the American Cancer Society RSG-22-022-01-CDP for support of this and other research work in ovarian cancer, and Dr. Deanna Teoh, Dr. Melissa Geller, and Dr. Rachel Vogel for helpful discussion on this topic. Dr. Martina Bazzaro would like to thank Dr. Simona Vertuani, Dr. Emma Hernlund, and Dr. Timothy Starr for the helpful discussion.

## Authors’ contributions

**EL**: conceptualization, data curation, funding acquisition, project administration, investigation, validation, writing of original draft. **VC**: data curation, formal analysis, investigation, methodology, visualization, writing of original draft. **MG**, data curation, methodology and investigation. **AX**: data curation, methodology, formal analysis, investigation, writing of original draft. **AN**: data curation, methodology, formal analysis, investigation. **FB**: data curation and methodology, **AS**: data curation, methodology and formal analysis. **SK**: data curation, methodology, formal analysis, investigation. **MB**: conceptualization, formal analysis, funding acquisition, project administration, investigation, resources, supervision, validation, writing of original manuscript.

## Ethics approval and consent to participate

Patients provided informed written consent for use of surgical tissue specimens for research. Archival tissues were used with approval from the Internal Ethics Review Board (Ethikkommission) of the Medical Faculty of the University of Tübingen (Project-Nr: 645/2012 BO2). The study was performed in accordance with the Declaration of Helsinki.

## Consent for publication

This manuscript does not contain individual details, images, videos, or other information that could identify specific individual patients.

## Data availability

The hyperlink/s to the publicly available diagnostic slides from the TCGA cohort (see Material and Methods) is provided <u>here</u>. Additional datasets used and/or analysed during the current study are available from the corresponding author upon reasonable request.

---

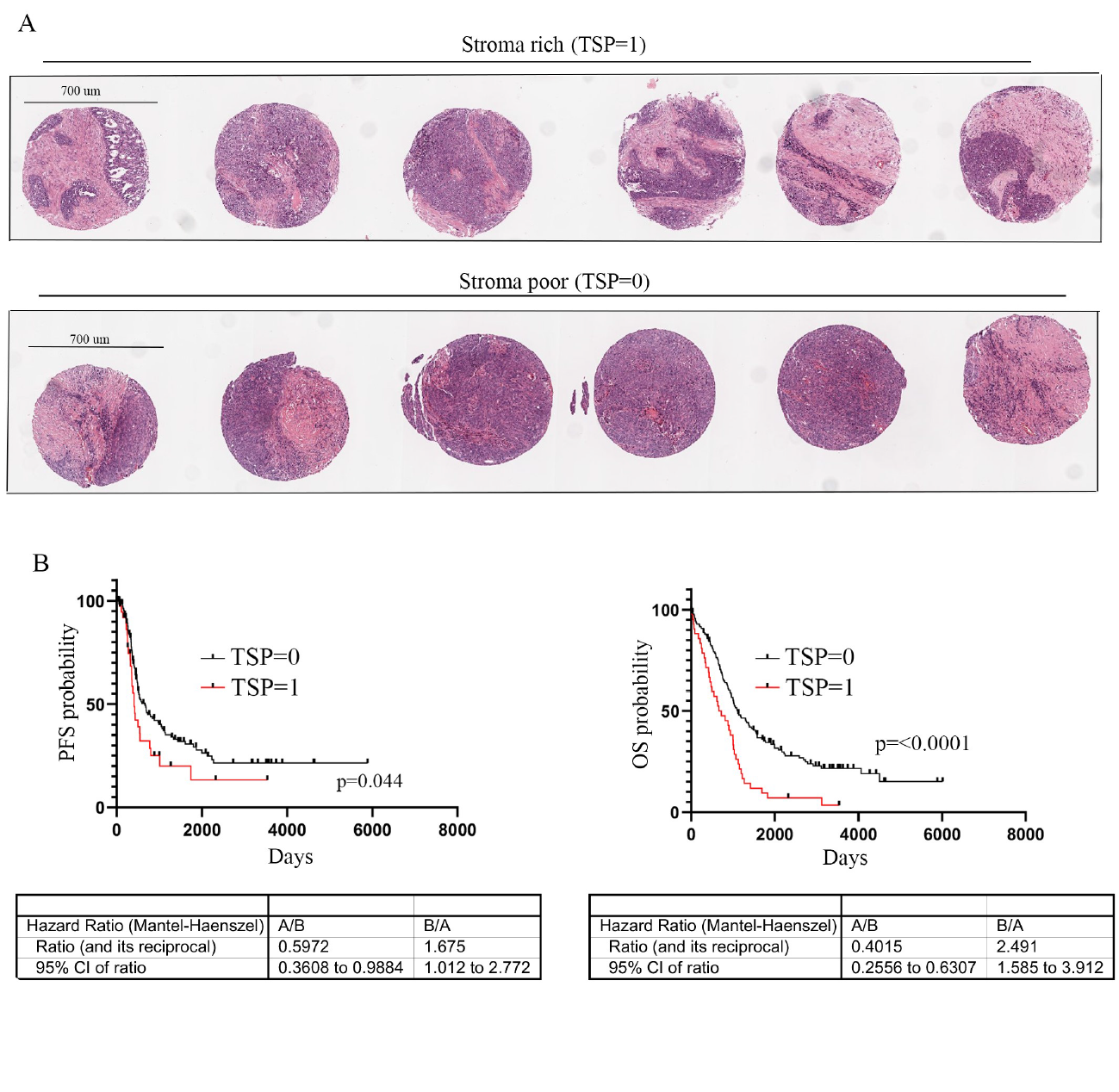

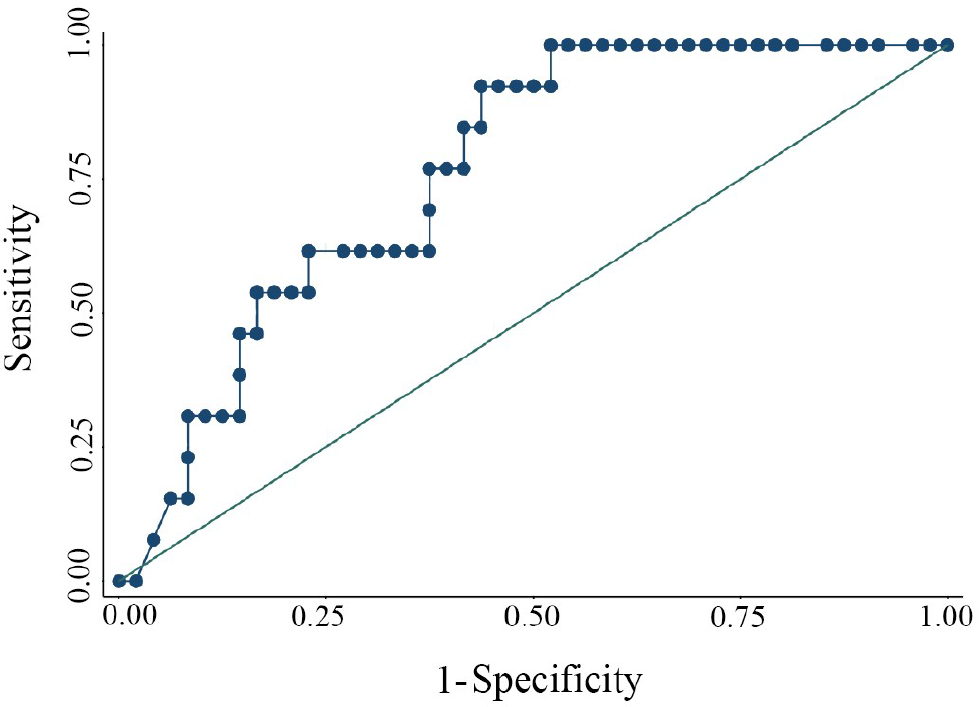

