## Supplementary Table 1 for "Clinical Significance of the Stromatic Component in Ovarian Cancer: Quantity Over Quality in Outcome Prediction"

**Supplementary Table 1**
**Patient Demographics and Clinical Characteristics of 192 Women Diagnosed with High Grade Serous Carcinoma (HGSC) of the Ovaries (University of Tubingen Cohort).** SD=standard deviation; FIGO=Federation Internationale de Gynecologie et d'Obstetrique.

| Characteristic | Level | Value |
|---|---|---|
| **Number of patients** | | 192 |
| **Age at diagnosis, mean (SD)** | | 63.7 (11.1) (n=192) |
| **Histology** | HGSC | 192 (100.0%) |
| **FIGO Stage** | 1 | 8 (4.2%) |
| | 2 | 14 (7.3%) |
| | 3 | 134 (69.8%) |
| | 4 | 36 (18.8%) |
| **Tumor (T)** | 1 | 9 (4.7%) |
| | 2 | 47 (24.5%) |
| | 3 | 130 (67.7%) |
| | Unknown | 6 (3.1%) |
| **Nodes (N)** | N0 | 75 (39.1%) |
| | N1 | 95 (49.5%) |
| | Unknown | 22 (11.5%) |
| **Metastases (M)** | 0 | 132 (68.8%) |
| | 1 | 36 (18.8%) |
| | Unknown | 24 (12.5%) |
| **Residual Disease** | No | 80 (41.7%) |
| | Yes | 108 (56.3%) |
| | Unknown | 4 (2.1%) |
