## Supplementary Table 2 for "Clinical Significance of the Stromatic Component in Ovarian Cancer: Quantity Over Quality in Outcome Prediction"

**Supplementary Table 2.**
**Multivariate analysis of progression free survival**

| Variable | Haz. ratio | P-value | [95% conf. | interval] |
|---|---|---|---|---|
| **TSP** | 1.586 | **0.015** | 1.093 | 2.302 |
| **Age** | 1.010 | 0.190 | 0.995 | 1.024 |
| **Metastasis** | 1.138 | 0.529 | 0.761 | 1.703 |
| **Residual Disease** | 2.038 | <0.001 | 1.436 | 2.892 |
